# A complex acoustical environment is necessary for maintenance and development in the zebra finch auditory pallium

**DOI:** 10.1101/2025.05.22.655494

**Authors:** Samantha M Moseley, C Daniel Meliza

## Abstract

Postnatal experience is critical to auditory development in vertebrates. The zebra finch (*Taeniopygia castanotis*) provides a valuable model for understanding how complex social-acoustical environments influence development of the neural circuits that support perception of vocal communication signals. We previously showed that zebra finches raised in the rich acoustical environment of a breeding colony (colony-reared, CR) perform twice as well in an operant discrimination task as birds raised with only their families (pair-reared, PR), and we identified deficits in functional properties within the auditory pallium of PR birds that could explain this behavioral difference. Here, using single-unit extracellular recordings from the L3 subdivision of field L and caudomedial nidopallium (NCM) at three developmental timepoints (18–20, 30–35, and 90–110 days post hatch), we tracked how experience affects the emergence of these functional properties. Whereas CR birds showed stable single-unit response properties from fledging to adulthood alongside improvements in population-level encoding, PR birds exhibited progressive deterioration in neural function. Deficits in PR birds began emerging at 18 days for population metrics and by 30 days for single-unit properties, worsening into adulthood. These included altered spike waveforms, firing rates, selectivity, discriminability, coding efficiency, and noise invariance. Notably, these deficits occurred despite PR birds receiving normal exposure to the song of a male tutor, suggesting that learning to sing is robust enough to compensate for impaired auditory processing. Our findings demonstrate that a complex acoustical environment is necessary for both maintenance and development of the cortical-level auditory circuits that decode conspecific vocalizations.

## Introduction

Many aspects of postnatal development in the vertebrate auditory system depend on experience. Altering an animal’s early environment by shifting the timing, feature distribution, or dynamics of auditory inputs produces long-lasting effects on intrinsic membrane properties (Kotak et al., 2005), synaptic connectivity (Dorrn et al., 2010; Sun et al., 2010), functional response properties (Zhang et al., 2001; de Villers-Sidani et al., 2007; Zhou and Merzenich, 2008; Insanally et al., 2009; Bao et al., 2013), and perceptually guided behavior (Han et al., 2007; Kover et al., 2013; Homma et al., 2020).

For humans, a particularly important instance of this plasticity occurs during the development of speech perception. Out of the distribution of speech sounds experienced during infancy, the brain forms a perceptual map for acoustical-articulatory categories (phonemes) integral to subsequent stages of language development (Werker and Lalonde, 1988; Kuhl et al., 1992; Ziegler et al., 2005). It is likely that this process of phonetic learning involves the same cellular and synaptic mechanisms underlying experience-dependent development in nonhuman auditory systems, but it remains unknown how these mechanisms are orchestrated to support rapid, reliable categorical decoding of high-dimensional, dynamic vocal signals like speech.

The zebra finch is a social songbird that breeds in dense colonies (Zann, 1996) and communicates individual identity and sexual fitness through a complex, learned vocalization called song (Elie and Theunissen, 2018). Zebra finches can reliably identify songs of other individuals (Yu et al., 2020) even when they are embedded in colony-like background noise (Narayan et al., 2007; Schneider and Woolley, 2013). An individual bird’s perception of song depends on its auditory experience during development (Clayton, 1990; Chen et al., 2017; Sturdy et al., 2001; Campbell and Hauber, 2009), and there are concomitant changes in the biophysical and functional properties of neurons in the auditory pallium (Phan et al., 2006; Amin et al., 2013; Moore and Woolley, 2019; Kudo et al., 2020), the avian homolog of cortex (Wang et al., 2010), that are consistent with the involvement of these areas in auditory memory and perception (Mello et al., 1995; Chew et al., 1996; Canopoli et al., 2014; Macedo-Lima and Remage-Healey, 2020; Yu et al., 2023).

Much of the research on developmental plasticity in the auditory pallium has focused on manipulating the song of an adult tutor (Adret et al., 2012; Yanagihara and Yazaki-Sugiyama, 2016), because for males these areas are also involved in acquiring a template from the tutor and providing feedback for sensorimotor learning and maintenance of song production (Bolhuis et al., 2000; London and Clayton, 2008). Thus, it has remained unclear which aspects of experience-dependent plasticity in the auditory pallium are related to song production, perception, or both. In our previous work, we developed an experimental design that allows us to separate the effects of the broader acoustical environment from effects specific to song production, by comparing zebra finches raised in the complex acoustical environment of a breeding colony (CR: colony-reared) to birds raised with just their families in acoustical isolation from the colony (PR: pair-reared). Males raised in both of these conditions develop normal song (Tchernichovski and Nottebohm, 1998), but both sexes show large differences in auditory perception and auditory pallial circuits (Chen and Meliza, 2020; Lu et al., 2023). Most recently, we found that CR adults perform twice as well in operant discrimination tasks compared to PR birds (Moseley and Meliza, 2025). Consistent with a deficit in auditory perception, neurons in the auditory pallium of PR adults exhibit lower firing rates, selectivity and discriminability, lower coding efficiency, less invariance to noise, and more redundancy compared to CR birds. It remains to be seen when and how these functional differences develop.

In the present study we sought to determine when the functional effects of a complex or impoverished acoustical environment occur relative to key milestones in zebra finch development (Doupe and Kuhl, 1999). Several previous studies have also examined the effects of experience on electrophysiological auditory responses at various time points in development (Amin et al., 2007; Miller-Sims and Bottjer, 2014; Schroeder and Remage-Healey, 2021, 2024), but using paradigms that manipulated the foreground tutor song inseparably from the acoustical background. Here, we performed single-unit extracellular recordings from male and female, CR and PR zebra finches at three time points: 18–20 days post hatch (dph), just after fledging, when auditory brainstem responses are mature (Amin et al., 2007) and birds are at the onset of the sensitive period for song memorization (Gobes et al., 2017); 30–35 dph, at the peak of the sensitive period and the age when we observe the largest effect of acoustical environment on intrinsic dynamics (Chen and Meliza, 2020); and 90–110 dph, when zebra finches reach sexual maturity and song has crystallized. We focused on the L3 subdivision of field L and the caudomedial nidopallium (NCM), two secondary pallial areas (Wang et al., 2010; Calabrese and Woolley, 2015), because these areas show the largest effect of the acoustical environment in adults. The results reported here include data from adults in L3/NCM that were part of the previous study (Moseley and Meliza, 2025). Hypothesizing that a complex social-acoustical environment is necessary for normal development of perception, we predicted that functional deficits would begin to emerge in PR birds during the sensitive period for song memorization and continue throughout development.

## Materials and Methods

### Animals

All procedures were performed according to National Institutes of Health guidelines and protocols approved by the University of Virginia Institutional Animal Care and Use committee. Male and female Australian zebra finches (*Taeniopygia castanotis*, recently recognized as a distinct species from the Sunda zebra finch; Olsson and Alström 2020) were bred in our local colony from 10 different pairs. All birds received finch seed (Abba Products, Hillside, NJ) and water ad libitum and were kept on a 16:8 light-dark schedule in temperature-and humidity-controlled rooms (22–24 °C).

Sex was determined from plumage coloration or by PCR amplification of the CHD-1 gene, which has different lengths on the Z and W sex chromosomes. Genomic DNA was isolated from feathers (Qiagen DNAeasy Blood and Tissue Kit) or from blood stored in a 5% Chelex-100 solution. The forward primer sequence was YTKCCAAGRATGAGAAACTG, and the reverse primer sequence was TCTGCATCACTAAAKCCTTT. These primers yield two products approximately 350 and 400 bp in length in females, the heterogametic sex in birds, whereas male DNA only yields the shorter product. No-template and positive controls were included in each PCR batch.

### Experimental Rearing Conditions

Zebra finches were reared in individual cages that were initially placed in the colony room, which housed around 70 male and female finches of varying ages at the time of the study. In the colony-reared (CR) condition, families remained in the colony room. In the pair-reared (PR) condition, families were moved to an acoustic isolation chamber (Eckel Industries) seven days after the first chick hatched. Hatchlings were fed by hand after the move as needed. Birds remained with their parents until they were used in an experiment or the youngest bird in the clutch was 35 dph, at which point the parents were removed. CR birds remained in the colony with other juveniles or in large single-sex flight aviaries. PR birds remained with their siblings in the acoustic isolation chamber.

### Surgery

Birds were surgically prepared for acute electrophysiology using previously described methods (Moseley and Meliza, 2025). The procedure was modified for birds recorded between 18–20 dph to accommodate the thinner, single-layer skull and to avoid having a recovery period while the birds were still under the care of their parents. On the day of the recording, juveniles were given three 0.03 mL intramuscular injections of 20% urethane separated by 20 min, then induced to a surgical anesthesia plane with 3.5% isoflurane (Southmedic). Feathers were removed from the top of the head and the bird was placed in a stereotaxic holder. The scalp was prepared with 7.5% povidone iodine (McKesson), Neosporin (Johnson & Johnson), and 2% Lidocaine (Henry Schein) prior to incision along the midline. After retracting the scalp, a metal pin was affixed to the skull 3 mm rostral to the y-sinus with Metabond (Parkell), and the skull over the recording site (approximately 3mm posterior of the y-sinus and 1.5 mm lateral of the midline) was removed with a scalpel. Birds were immediately transferred to an acoustically isolated recording chamber (Eckel Industries), where they were restrained in a padded 50 mL conical tube and the head pin was attached to a stand. The dura was thinned using an Ultra Needle (Electron Microscopy Sciences), a well was formed with Kwik-Cast around the recording site, and the ground wire was placed in the well.

### Data Acquisition

Neural activity was recorded from the adults and one P30 bird using 128-channel two-shank silicon probes (128 AxN Sharp, Masmanidis Lab). After this probe was discontinued, we switched to 64-channel two-shank silicon electrodes (ASSY-77 H6, Cambridge NeuroTech) for the remaining P30 and P18 birds. Extracellular data were filtered between 1–9987 Hz and digitized at 30 kHz using an Open Ephys Acquisition Board connected to a computer running Open Ephys GUI software (v0.5.3.1). The electrode was coated with DiI (ThermoFisher Scientific) and inserted at a dorso-caudal to ventro-rostral angle that confined the penetrations to the deep/secondary areas L3 and NCM (caudomedial nidopallium). After contacting the brain surface, the probe was advanced at 1 µm/s. A collection of zebra finch songs not used in the main study was played during electrode advancement to monitor auditory responsiveness. Electrode movement was stopped once the local field potentials and spiking indicated reliable auditory responses across the whole probe. Once the electrode was in position, the well was filled with phosphate-buffered saline (PBS; in mM: 10 Na2HPO4, 154 NaCl, pH 7.4). After responses to the full set of stimuli were recorded, we either withdrew the electrode and inserted it in a new location or terminated the recording.

### Stimuli

We used the same stimuli as in Moseley and Meliza (2025). Briefly, the stimuli comprised songs selected from recordings of 10 unfamiliar male conspecifics. Songs included 1–2 motifs and were 2.52 *±* 0.28s in duration (mean *±* SD; range 1.92–2.88s). The songs were filtered with a 4th-order Butterworth highpass filter with a cutoff of 150 Hz, a 2 ms squared cosine ramp was applied to the beginnings and the ends, and the waveforms were rescaled to have an RMS amplitude of −30 dB FS. The songs were then embedded in synthetic noise generated by the McDermott and Simoncelli (2011) algorithm to match the statistics of our colony nestbox recordings. The amplitude of the background was varied to give digital signal-to-noise ratios (SNRs) ranging from 70 to −10 dB. The foreground songs were presented at approximately 60 dB_A_ SPL, so the noise at the highest SNR level (–10 dB_A_ SPL) was well below the ambient noise level in the acoustic isolation boxes (around 30 dB_A_ SPL) and below the hearing threshold of the finches (Hashino and Okanoya, 1989). For simplicity, we use the SNR values of the digital sound files as labels.

Acoustic stimuli were presented with custom Python software (https://github.com/melizalab/open-ephys-audio; v0.1.7) through a Focusrite 2i2 analog-to-digital converter, a Samson Servo 120a amplifier, and a Behringer Monitor Speaker 1C. The gain on the amplifier was adjusted so that the foreground songs had an RMS amplitude of 60 dB_A_ SPL at the position of the bird’s head. The songs were combined into sequences, each consisting of 10 permutations following a balanced Latin square design (i.e., each song occurred once in each of the 10 positions and was preceded by a different song in each permutation). The songs within each sequence were separated by gaps of 0.5 s, and the sequences were presented with a gap of 5 s. The sequences were embedded in a background consisting of synthetic colony noise that began 2 s before the first song and ended 2 s after the end of the last song.

### Histology

Immediately after recording, animals were administered a lethal intramuscular injection of Euthasol (Vibrac) and perfused transcardially with 10 U/mL solution of sodium heparin in PBS followed by 4% formaldehyde in PBS. Brains were removed from the skull, postfixed overnight in 4% paraformaldehyde, blocked along the midline, and embedded in 3% agar (Bio-Rad). Sections were cut at 60 µm on a vibratome and mounted on slides. After drying for 30 min, the sections were coverslipped with Prolong Gold with DAPI (ThermoFisher, catalog P36934; RRID:SCR_05961). The sections were imaged using epifluorescence with DAPI and Texas Red filter cubes to locate the DiI-labeled penetrations. Because we only painted the electrode once at the beginning of the experiment and made sure to move it 100 µm or more between penetrations, we were able to reconstruct the locations of multiple recording sites per animal.

### Spike Sorting and Classification

Spikes were sorted offline with Kilosort 2.5. Clusters were automatically excluded if they had a contamination score higher than 10 or contained fewer than 100 spikes (corresponding to a rate of 0.029 Hz over the 58 min session), as it was not possible to assess quality with so few events. Clusters were further curated by visual inspection (phy v2.0; https://github.com/cortex-lab/phy) for spheroid distribution in the feature space, very low refractory period violations in the autocorrelogram, stable firing rate throughout the recording, and a typical average spike waveform. Individual spikes were excluded as artifacts if their amplitude was more than 6 times the amplitude of the average spike from that unit. To classify units as narrow-spiking (NS) or broad-spiking (BS), the spike waveforms for each unit were averaged after resampling to 150 kHz using sinc interpolation. Taking the time of the maximum negative deflection (trough) as 0, the spike width was defined as the time of the maximum positive deflection (peak) following the trough, and the peak/trough ratio was measured from the (absolute) magnitude of the peak and trough. A Gaussian mixture model with two groups was fit to these features, and units were classified as narrow-spiking if they were assigned to the group with the lower spike width and higher peak/trough ratio.

### Experimental Design and Statistical Analysis

The objective of the study was to characterize how the acoustical environment affects development of functional responses properties in the zebra finch auditory pallium. Electrophysiological recordings were collected from birds in three different age groups (P8: 18–20 dph; P30: 30–35 dph; P90: 90–110 dph). Table 1 gives the number of birds, recordings, and units in each condition. The data from 3 CR adults and 4 PR adults have been previously reported (Moseley and Meliza, 2025); one adult from each rearing condition was added for this study to better balance sex. The main independent variables in this study were rearing condition (CR or PR), age (P18, P30, or P90), and spike types (BS or NS). Birds were chosen from CR and PR clutches at random while maintaining a balance of males and females. Sex and family were not included as analysis variables due to the limited sample size. The experimenter was aware of the rearing condition of each animal. Unless otherwise noted, we included every well-isolated single unit in the analysis.

**Table 1.** Sample sizes for birds, recordings, units, and pairs of simultaneously recorded units. Each recording comprises responses to a complete stimulus set at a unique location in the auditory pallium. Units comprise all the well-isolated single units recorded in each condition, and pairs comprise all the simultaneously recorded units in each condition.

|  |  | P18 |  | P30 |  | P90 |  | Total |  |
| --- | --- | --- | --- | --- | --- | --- | --- | --- | --- |
|  |  | CR | PR | CR | PR | CR | PR | CR | PR |
| Birds |  | 2♂ 2♀ | 2♂ 2♀ | 2♂ 2♀ | 3♂ 2♀ | 2♂ 2♀ | 3♂ 2♀ | 6♂ 6♀ | 8♂ 6♀ |
| Recordings |  | 8 | 8 | 7 | 8 | 8 | 13 | 23 | 29 |
| Units | BS | 232 | 286 | 292 | 347 | 478 | 230 | 1002 | 863 |
|  | NS | 66 | 74 | 115 | 72 | 90 | 139 | 271 | 285 |
|  | Total | 298 | 360 | 407 | 419 | 568 | 369 | 1273 | 1148 |
| Pairs | BS | 470 | 1291 | 1231 | 3016 | 2012 | 420 | 3713 | 4727 |
|  | NS | 234 | 207 | 1272 | 387 | 387 | 236 | 1893 | 830 |
|  | Total | 704 | 1498 | 2503 | 3403 | 2399 | 656 | 5606 | 5557 |

Unless otherwise noted, data analysis procedures were the same as in our previous study (Moseley and Meliza, 2025) but are reported in full here for completeness. Parameters of generalized linear mixed effects models (GLMMs) were estimated in R using lme4 (version 1.1.35.3). Marginal effects, contrasts, and confidence intervals were computed from the models using emmeans (version 1.10.2).

Spike trains were separated into non-overlapping intervals corresponding to songs, beginning at the start of the song and ending 0.5 s after the end of the song. This gave us 10 trials per 10 songs at each of 11 SNR levels per unit. For spontaneous firing, we used 1 s intervals taken from the periods of silence at the beginning of each song sequence, for a total of 110 trials per unit.

Spontaneous and evoked firing rates were quantified using generalized linear mixed-effects models (GLMMs) with spike count as the dependent variable with a Poisson distribution. The model specification was `n_events ~ age_group*spike*rearing + offset(duration) + (1| unit) + (1|foreground)+ (1| bird) <, where age_group*spike*rearing` defines fixed effects for age groups, cell type, rearing condition, and their interactions; offset(duration) defines a constant offset term for the duration of the interval in which the spikes were counted; and (1|unit), (1|foreground), and (1|bird) define random intercepts for each unit, stimulus (omitted from the model for spontaneous rate), and bird. In contrast to our previous study, we were able to include a random effect for bird because the design was better balanced across age groups. Contrasts are reported on the log ratio (LR) scale.

Discriminability was calculated for each unit by computing the pairwise similarity between all of its song-evoked responses. Responses were truncated to the duration of the shortest stimuli to remove information encoded in the response length. One stimulus was excluded because it was much shorter than the others. We used the SPIKE-synchronization metric (Kreuz et al., 2015) implemented in the pyspike Python package (v0.7.0; http://mariomulansky.github.io/PySpike/), yielding a 90 *×* 90 matrix of similarity scores that were used to train a *k*-nearest neighbors classifier (scikit-learn v1.3.2). That is, each trial was classified by finding the *k* most similar trials (the neighbors) and assigning it the label that was the most common in that group. We used *k* = 9, but other values produced nearly identical results. This yielded a 9 *×* 9 confusion matrix with one dimension corresponding to the stimulus that was actually presented and the other to the label that was assigned to the trial. Each unit received a score for the proportion of trials that were correctly classified. We used a GLMM with the number of correctly classified trials as the binomial dependent variable to quantify the effect of rearing condition. The model specification in R was `cbind(n_correct, 90 - n_correct) ~ age_group*spike*rearing + (1|unit) + (1|bird)`. Contrasts are reported on the log odds ratio (LOR) scale. Exponentiating LOR gives the fold difference between groups or conditions in the odds of correctly decoding which stimulus was presented from a single unit’s firing patterns. Discriminability was also used to classify neurons as auditory: a neuron was deemed to be auditory if its average score for all the songs was more than 1.64 standard deviations (the one-sided 95% confidence level for a z-test) above the mean for 500 permutations in which the song labels were randomly shuffled.

Selectivity was calculated for each unit as 1 minus the proportion of songs that evoked a firing rate that was significantly greater than the spontaneous rate. We used a generalized linear model with spike count as the Poisson-distributed dependent variable to determine significance. For the rare neurons that did not spike at all during the spontaneous intervals (1.1%, *n* = 27/2394), a single spike was added to the spontaneous count to regularize the model estimates. To quantify the effects of age and rearing condition, we used a GLMM with the number of songs with rates above spontaneous as the binomial dependent variable. The model specification in R was `cbind(10 - n_responsive, n_responsive) ~ age_group*spike*rearing +(1|unit) + (1|bird)`. Note that defining responsive songs as “failures” gives a selectivity score where 0 is a unit that responds to all the stimuli and 1 is a unit that responds to none. Only auditory units were included in this analysis. Contrasts are reported on the LOR scale. Exponentiating LOR gives the fold difference in the odds that a neuron will not respond to a randomly chosen stimulus.

Signal correlations were computed using the number of spikes evoked in the interval between the start of each song and 0.5 s after its end. For each pair of simultaneously recorded neurons, the signal correlation was calculated as the Pearson correlation coefficient between the trial-averaged firing rates to the ten songs. Only BS–BS and NS-NS pairs were considered, due to the difficulty of interpreting BS–NS correlations. Only auditory units were considered, and pairs recorded on the same channel were excluded to limit the influence of spike-sorting errors. The effect of rearing condition and age on signal correlations was estimated using linear regression in R, with age group, cell type, rearing condition, and their interactions as predictors. We did not include random effects in this analysis because it was not clear how to consider repeated measures with pairs of neurons. Because we used extracellular probes with different physical arrangements of recording sites for different age groups and were concerned that this might impact the spatial sampling of simultaneously recorded pairs, we did not examine noise correlations.

Stimulus reconstruction from population responses was implemented using a linear decoder. The model is similar to the spectrotemporal receptive field, in which the expected firing rate of a single neuron at a given time point *t* is modeled as a linear function of the stimulus spectrogram immediately prior to *t*, but in the linear decoding model, the relationship is reversed, and the expected stimulus at time *t* is modeled as a linear function of the response that follows. Using a discrete time notation where *s*_*t*_ is the stimulus in the time bin around *t*, and *r*_*t*_ is the response of a single neuron in the same time bin, then the expected value of the stimulus is given by

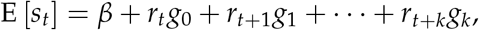

where *k* is the number of time bins one looks into the future, **g** = (*g*_0_, *g*_1_, …, *g*_*k*_) are the linear coefficients of the model, and *β* is a constant intercept term. If the errors are independent and normally distributed around the expectation with constant variance *σ*^2^, then this is an ordinary linear model. If there are *n* time bins in the stimulus, then the stimulus is a vector **s** = (*s*_0_, …, *s*_*n*_) drawn from a multivariate normal distribution. In vector notation,

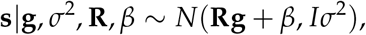

where **R** is the *n × k* Hankel matrix of the response. Without any loss of generality, the model can be expanded to include the responses of multiple neurons. If there are *p* neurons, then *r*_*t*_ becomes a *p*-element vector (*r*_1,*t*_, …, *r*_*p,t*_), **R** becomes a *n × pk* matrix formed by concatenating the Hankel matrices for each of the neurons, and **g** becomes a *pk*-element vector.

In this study, the response matrix was constructed from the peristimulus time histograms (PSTHs) averaged over 10 trials using a bin size of 2.5 ms. Because the number of units sampled varied by rearing condition and age, we used a bootstrapping approach to randomly select a fixed number of units without replacement from each condition and compared conditions using matched ensemble sizes. We used 100 replicates from each condition except when the ensemble size was the same as the number of units for that condition. We used a *k* of 80, corresponding to lags from 0–200 ms. The response matrix was projected into a basis set consisting of 20 nonlinearly spaced raised cosines (Pillow et al., 2005). The width of each basis function increased with lag, giving higher temporal resolution at short lags and lower resolution at long lags. This allowed the inclusion of longer lags without exploding the number of parameters.

Stimuli were converted to time-frequency representations using a gammatone filter bank, implemented in the Python package gammatone (version 1.0) with 40 log-spaced frequency bands from 1–8.5 kHz, a window size of 5 ms, and a step size of 2.5 ms. Power was log-transformed.

We used ridge regression to estimate the parameters of the model because the number of parameters *pk* was typically larger than the number of time bins *n*. The decoder performance was estimated using a 10-fold cross-validation strategy. There were 10 data folds generated by holding out each of the songs. For each fold, 9 songs at the highest SNR (70 dB) were used to estimate *ĝ*_*λ*_, where *λ* denotes the ridge shrinkage penalty. We tested 30 *λ* values on a logarithmic scale from 10^−1^ – 10^7^. *ĝ*_*λ*_ was used to decode the stimulus 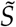 from the responses to the held-out song 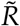, also at 70 dB SNR. The score for a given prediction was quantified as the adjusted coefficient of determination *R*^2^. The cross-validated prediction score was computed by averaging scores across folds for each value of *λ* and then taking the best score. To estimate noise invariance, we used the trained model to decode the stimulus from the responses to the held-out song at the full range of SNR levels. Ridge regression and cross-validation were performed in Python with the scikit-learn library (v1.3.2).

We introduced a new noise invariance metric in this study that was calculated for individual units using spike train similarity. Recall that we presented 10 different permutations of 10 songs against a constant background at 11 different SNR levels. For each permutation, we compared the response at each SNR between 35 and −10 dB SNR to the response at the lowest noise level (70 dB SNR) to quantify how much the firing pattern was affected by increasing levels of noise, averaging across permutations to obtain *S*_*F*_(*SNR*), the similarity to foreground. For each noise level, we calculated the average pairwise similarity between different permutations to quantify how much the firing pattern was driven by the constant background, *S*_*B*_(*SNR*). Noise invariance was calculated as *S*_*F*_(*SNR*) − *S*_*B*_(*SNR*). This measure is based on the same principle as the extraction index employed by Schneider and Woolley (2013), but using a spike train metric of response similarity instead of correlation coefficient. We used a linear mixed-effects model with invariance as the dependent variable to quantify the effect of rearing condition, spike type, and age group across SNR. The model specification in R was `invariance ~ SNR*rearing*age_group*spike + (1|unit) + (1|bird).`

## Data and Code Availability

Data were analyzed using custom Python and R code (https://github.com/melizalab/cr-pr-juveniles, to be released after initial peer review). Stimuli and spike time data have been deposited on figshare (url TBD, to be released after initial peer review).

## Results

### An impoverished acoustical environment dampens neural activity

To test how the acoustical environment impacts the development of functional response properties in L3 and NCM, we compared birds raised in our breeding colony (colony-reared: CR) to birds raised in acoustical isolation with parents and siblings (pair-reared: PR) at 18–20 dph (P18; *n* = 4 CR, 4 PR), 30–35 dph (P30; *n* = 4 CR, 5 PR), and 90–110 dph (P90; *n* = 4 CR, 5 PR). We recorded extracellular responses under urethane anesthesia to 10 unfamiliar conspecific songs embedded in varying levels of synthetic colony noise, confirming electrode placement post hoc (Fig. 1A). In all experimental conditions, single unit waveforms clustered into broad-spiking (BS) and narrow-spiking (NS) groups (Fig. 1B), but there were subtle differences in the distribution of spike shapes within these categories across age groups and acoustical environments (Fig. 1C). For simplicity and consistency with prior work, we used a global Gaussian mixture model to classify units from all conditions as BS or NS. Average spike width and peak-to-trough ratio remained stable from P18 to P90 for both cell types in the CR birds but not in the PR birds (Fig. 1C,D). Compared to BS units from age-matched CR birds, the BS units in PR birds had narrower spikes at P90 (Fig. 1D) and shallower rebounds (lower peak-to-trough ratio; Fig. 1B) at P30 (Fig. 1E).

**Figure 1.**
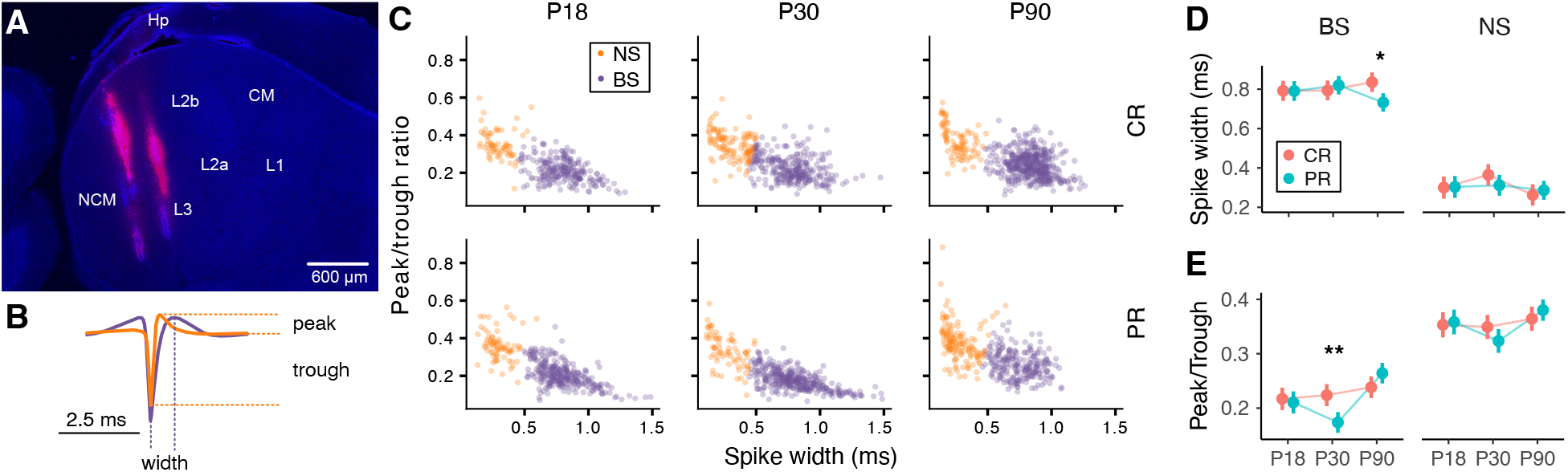
Recording methods and extracellular spike waveforms. (**A**) Example fluorescence image of a parasagittal section of zebra finch auditory pallium. Blue is DAPI and red is DiI that was coated onto the electrode. Each track is a separate penetration. The second electrode shank is found in more lateral section. (**B**) Average waveforms for all broad-spiking (BS, purple) and narrow-spiking (NS, orange) units in the study. Dashed lines illustrate measurement of waveform features used to classify individual units. (**C**) Scatter plots of peak-to-trough ratio and spike width for individual units in each age group and rearing condition. Units were classified as BS or NS using a Gaussian mixture model for these features, fit to all the units. There was a statistically significant interaction effect between age group and rearing condition on spike shape features within BS and NS (MANOVA: Wilks’ *λ* = 0.34, *F*_12,4816_ = 290, *p <* 0.001). (**D**) Average spike width in each age group and rearing condition for units classifed as BS (left) and NS (right). Filled circles show means for colony-reared (salmon) and pair-reared (cyan) conditions with 90% credible intervals for the whiskers (GLMM: see Materials and Methods). Spikes in BS units were wider in adult CR birds compared to adult PR birds (*β* = 0.10 *±* 0.04 ms, *t*_20.4_ = 2.6, *p* = 0.017). (**E**) Average peak-to-trough ratio in each age group and rearing condition. At P30, spikes in BS units had deeper troughs in CR birds compared to PR birds (*β* = 0.050 *±* 0.016, *t*_20.9_ = 3.1, *p* = 0.005). Within BS cells from the PR condition, there were significant differences between P90 and P18 (*β* = 0.053 *±* 0.016, *t*_21.4_ = 3.3, *p* = 0.008) and between P90 and P30 (*β* = 0.091 *±* 0.016, *t*_22.7_ = −5.8, *p <* 0.001). Within NS cells from the PR condition, there was a significant difference between P90 and P30 (*β* = 0.057 *±* 0.017, *t*_35.5_ = 3.3, *p* = 0.007).

Our initial analysis focused on responses to stimuli where the background noise was inaudible (70 dB SNR). At all ages, NS neurons had higher firing rates and responded to a greater proportion of stimuli than BS neurons (Fig. 2). Average spontaneous and evoked firing rates of NS and BS cells depended on age and experience. In the CR condition, there was no significant change in firing rates across P18, P30, and P90 (Fig. 3A,B). Evoked firing rates for BS and NS neurons were elevated in PR birds at P30, relative to age-matched CR birds. Spontaneous and evoked firing rates of the NS units in PR birds both decreased between P30 and P90, leading to a 5-fold difference between CR and PR adults in NS spontaneous rates and a 3.6-fold difference in NS evoked rates. Across age groups, we sampled more BS units compared to NS units (Fig. 3C). The proportion of BS units sampled remained consistent across development for CR birds, but there was a significant decrease between P30 and P90 in PR adults.

**Figure 2.**
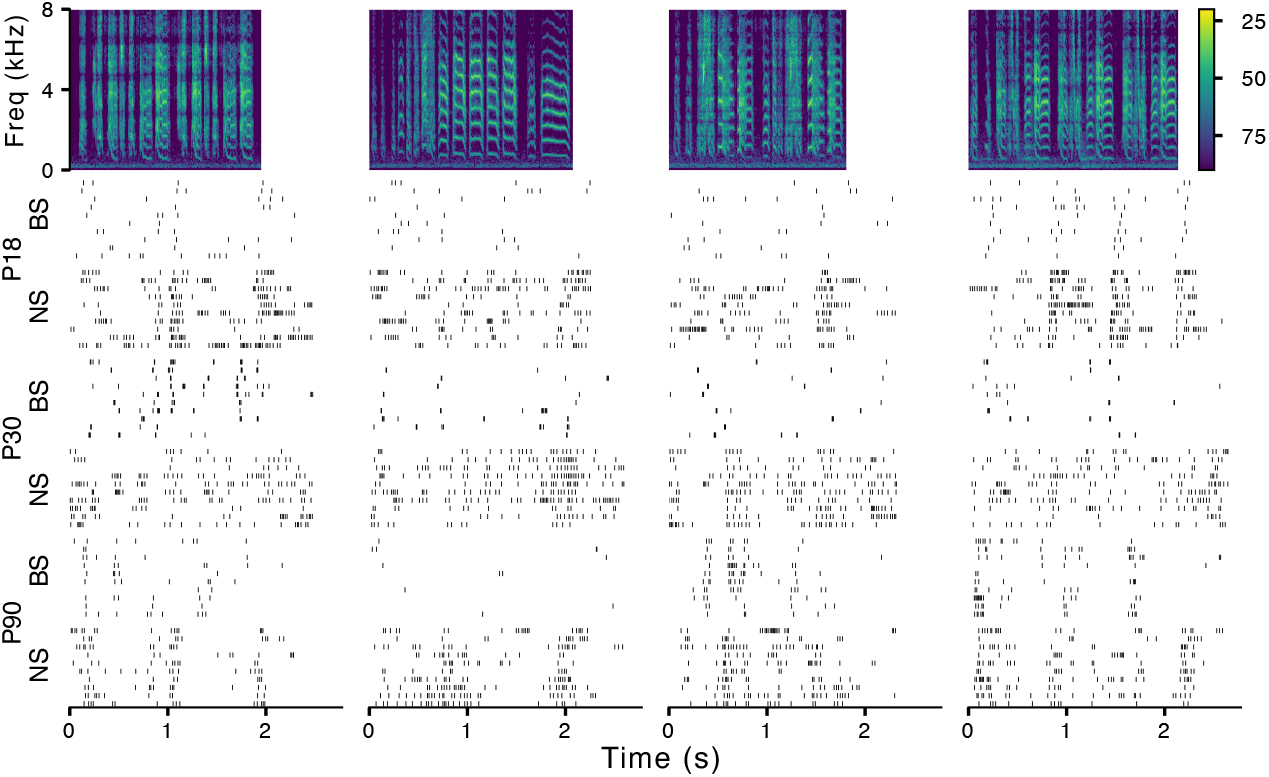
Example auditory responses in L3/NCM. Raster plots of responses from example NS and BS example units from CR birds in each age group to four unfamiliar zebra finch songs (spectrograms, top) across the three developmental time points. Spectrogram color indicates amplitude relative to the full scale of the sound file (dB FS).

**Figure 3.**
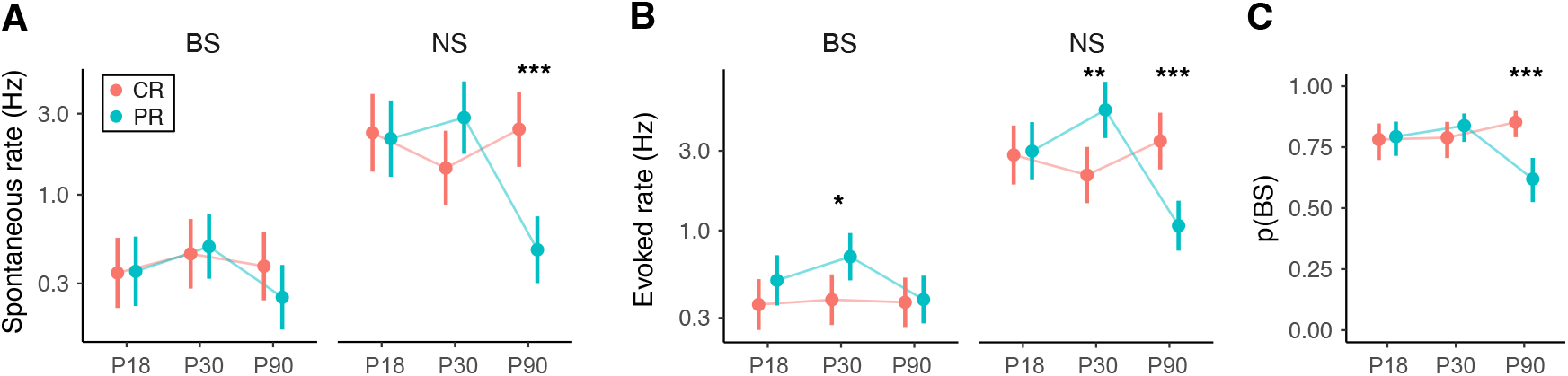
Effects of acoustical environment on firing rates. (**A**) Average spontaneous firing rate across rearing conditions and age groups. Filled circles represent marginal means for colony-reared (salmon) and pair-reared (cyan) conditions with 90% credible intervals for the whiskers (GLMM; see Materials and Methods). Asterisks represent a significant post-hoc difference between rearing conditions within each age group and cell type (*: *p <* 0.05; **: *p <* 0.01, ***: *p <* 0.001). In adults, spontaneous rates of NS units were lower in PR adults compared to CR adults (log ratio [LR] = −1.64 *±* 0.41; *z* = −3.95; *p <* 0.001). Within NS cells from the PR condition, the spontaneous rate at P90 was lower than at P18 (LR = −1.51 *±*0.42; *z* = −3.60; *p <* 0.001) and at P30 (LR =−1.79 *±* 0.41; *z* = −4.42; *p <* 0.001). (**B**) Average evoked firing rates (stimuli with inaudible background only). In the P30 group, evoked rates were elevated in PR compared to CR birds for BS units (LR =−0.59 *±* 0.29; *z* = −2.05; *p* = 0.04) and NS units (LR = 0.90*±*0.33; *z* = −2.71; *p* = 0.006). In adults, evoked rates were lower in PR compared to CR birds in the NS units only (LR = −1.17 *±* 0.32; *z* = −3.70; *p <* 0.001). Within NS cells from the PR condition, the average evoked rate at P90 was less than at P18 (LR = −1.02 *±* 0.32; *z* = −3.21; *p* = 0.004) and at P30 (LR = −1.59 *±* 0.31; *z* = −5.08; *p <* 0.001). (**C**) Proportion of neurons at each site with broad spike waveforms. There were proportionately fewer BS neurons sampled in PR adults compared to CR adults (log odds ratio [LOR] = −1.26 *±* 0.35; *z* = −3.6; *p <* 0.001) and compared to PR chicks at P18 (LOR = −0.85 *±* 0.35; *z* = −2.44; *p* = 0.04) and at P30 (LOR = −1.16 *±* 0.35; *z* = −3.30; *p* = 0.003

### Development of discriminability, selectivity, and signal correlation

To determine if the effects of the acoustical environment on the development of spontaneous rate and overall responsiveness were reflected in the functional properties of L3/NCM neurons, we measured discriminability, selectivity, and signal correlations in NS and BS cells from each age group and rearing condition using the same methods we employed previously (Moseley and Meliza, 2025).

Discriminability quantifies how reliably and distinctively a unit responds to different stimuli using a spike-train similarity metric to compare the response in each trial to every other trial. As illustrated in Figure 4A–B, NS neurons tended to respond to most songs with distinctive firing patterns (Fig. 2), which was reflected in high similarity among trials for the same song and low similarity among trials from different songs. This resulted in high performance for a k-neighbors classifier trained to decode which stimulus was presented. BS neurons tended to respond to a smaller proportion of songs, which resulted in high within-song similarity for those songs only and higher similarity with trials for different songs, and the classifier could only accurately decode a few of the stimuli (Fig. 4A–B, bottom). Conversely, selectivity measures the proportion of stimuli that evoke a significant increase in firing rate. As illustrated in Figure 4C, NS neurons tended to be less selective than BS neurons.

**Figure 4.**
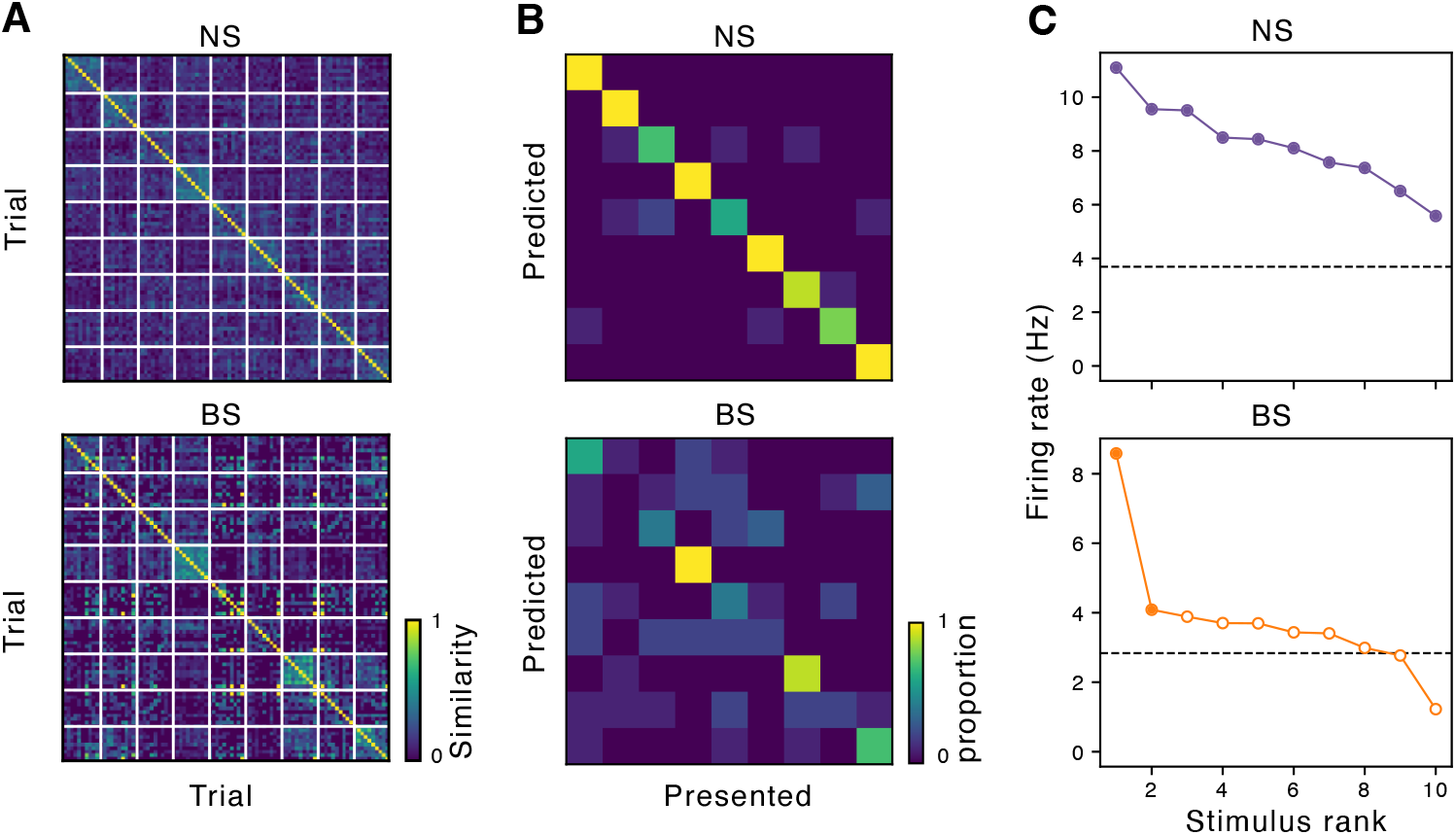
Examples of single-unit metrics of discriminability and selectivity. (**A**) Spike train similarity between all pairs of trials (inaudible noise stimuli only) for example BS and NS units. Rows and columns are sorted by song, indicated by grid of white lines. (**B**) Confusion matrices from predictions of a classifier using spike train similarities in *A*. The diagonal corresponds to correct classifications. (**C**) Average firing rates of the example units in A,B evoked by the songs, sorted in rank order. Dashed line represents the spontaneous rate. Filled circles represent firing rates significantly greater than the unit’s spontaneous firing rate (GLM; see Materials and Methods).

Consistent with the examples in Figure 4A–B, discriminability, quantified as the proportion correct of the classifier’s performance for each unit, was higher for NS neurons compared to BS neurons across all ages and rearing conditions (Fig. 5A; LOR = 1.34 *±* 0.05; *z* = 27.2; *p <* 0.001). Discriminability remained at the same low level across P18, P30, and P90 for BS neurons in both rearing conditions. In NS neurons, discriminability increased with age for CR birds but decreased in PR birds, leading to a more than 2.5-fold difference between the groups at P90.

**Figure 5.**
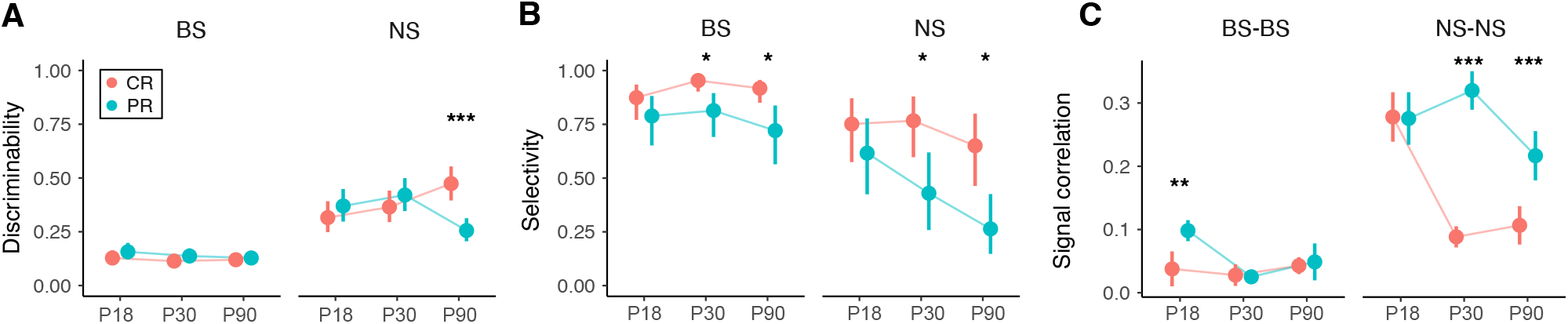
Experience-dependent development of discriminability, selectivity, and signal correlation. (**A**) Average discriminability, measured as the proportion of correctly decoded stimuli, for each cell type, rearing condition, and age group, same format as in Figure 3 In adults, discriminability of NS units was lower in PR birds compared to CR birds (LOR = −0.96 *±* 0.26; *z* = −3.7; *p <* 0.001). Within NS units from PR birds, discriminability decreased between P30 and P90 (LOR = −0.75 *±* 0.26; *z* = −2.91; *p* = 0.01). Within NS units from CR birds, discriminability increased between P18 and P90 (LOR = 0.66 *±* 0.24; *z* = 2.38; *p* = 0.046). (**B**) Average selectivity, measured as the proportion of stimuli evoking responses greater than the spontaneous rate. Selectivity for BS units was lower in PR birds compared to CR birds at P30 (LOR = −1.55 *±* 0.63; *z* = −2.46; *p* = 0.01) and P90 (LOR =−1.46 *±* 0.59; *z* = −2.50; *p* = 0.01) and for NS units at the same ages (P30: LOR = −1.47 *±* 0.67; *z* = −2.19; *p* = 0.028; P90: LOR = −1.65*±*0.64; *z* = −2.57; *p* = 0.01). (**C**) Signal correlations between simultaneously recorded BS-BS and NS-NS pairs. For pairs of BS neurons, signal correlations were higher in PR birds compared to CR birds at P18 (*β* = 0.06 *±* 0.02; *t*_11151_ = 3.09, *p* = 0.002). For pairs of NS neurons, signal correlations were higher in PR birds compared to CR birds at P30 (*β* = 0.23 *±* 0.02; *t*_11151_ = 11.0; *p <* 0.001) and P90 (*β* = 0.11 *±* 0.03; *t*_11151_ = 3.67; *p <* 0.001).

Across all ages and rearing conditions, selectivity was higher on average for BS neurons compared to NS neurons (Fig. 5B; LOR = 1.51 *±* 0.16; *z* = 9.2; *p <* 0.001). There was no statistically significant change over development in selectivity within BS or NS neurons for either rearing condition, but selectivity was lower in both cell types at P30 and P90 in PR birds compared to age-matched CR birds.

We also examined how signal correlations between simultaneously recorded pairs of BS and NS neurons changed over development. Higher signal correlations indicate that neurons are tuned to similar stimuli, leading to greater redundancy and less information about the stimulus in the population. Across all ages and rearing conditions, BS–BS pairs were less correlated than NS–NS pairs (Fig. 5C; *β* = −0.16 *±* 0.01; *t*_11151_ = −17.1; *p <* 0.001). BS–BS pairs were more correlated in P18 PR birds compared to CR birds at the same age; this difference disappeared at P30 and P90. NS–NS pairs showed the opposite trend, with no difference between CR and PR birds at P18 but more correlated in PR birds compared to CR birds at P30 and P90. There was a large decrease in signal correlation in CR birds between P18 and P30 that did not occur in the PR birds.

### Development of linear decodability and noise invariance

In our previous study of the auditory pallium in adults, we found that the spectrotemporal structure of song can be decoded from smaller populations of neurons in CR birds compared to PR birds and that this information is more invariant in the presence of colony-like background noise (Moseley and Meliza, 2025). To test how this difference emerges over development, we applied the same linear decoder analysis using responses to stimuli embedded in synthetic colony noise at signal-to-noise ratios (SNRs) between 70 and −10 dB. In all age groups and rearing conditions, firing patterns of individual neurons became less consistent as the noise level increased (Fig. 6A). A linear decoder was trained on the trial-averaged response of the population to 9 of the 10 songs at the highest SNR and then used to decode the stimulus from the response to the remaining song across all SNR levels (Fig. 6B). Spectrograms decoded from the responses to stimuli with high SNRs maintained many features of the original clean song, but the similarity decreased as the noise level was increased (Fig. 6B).

**Figure 6.**
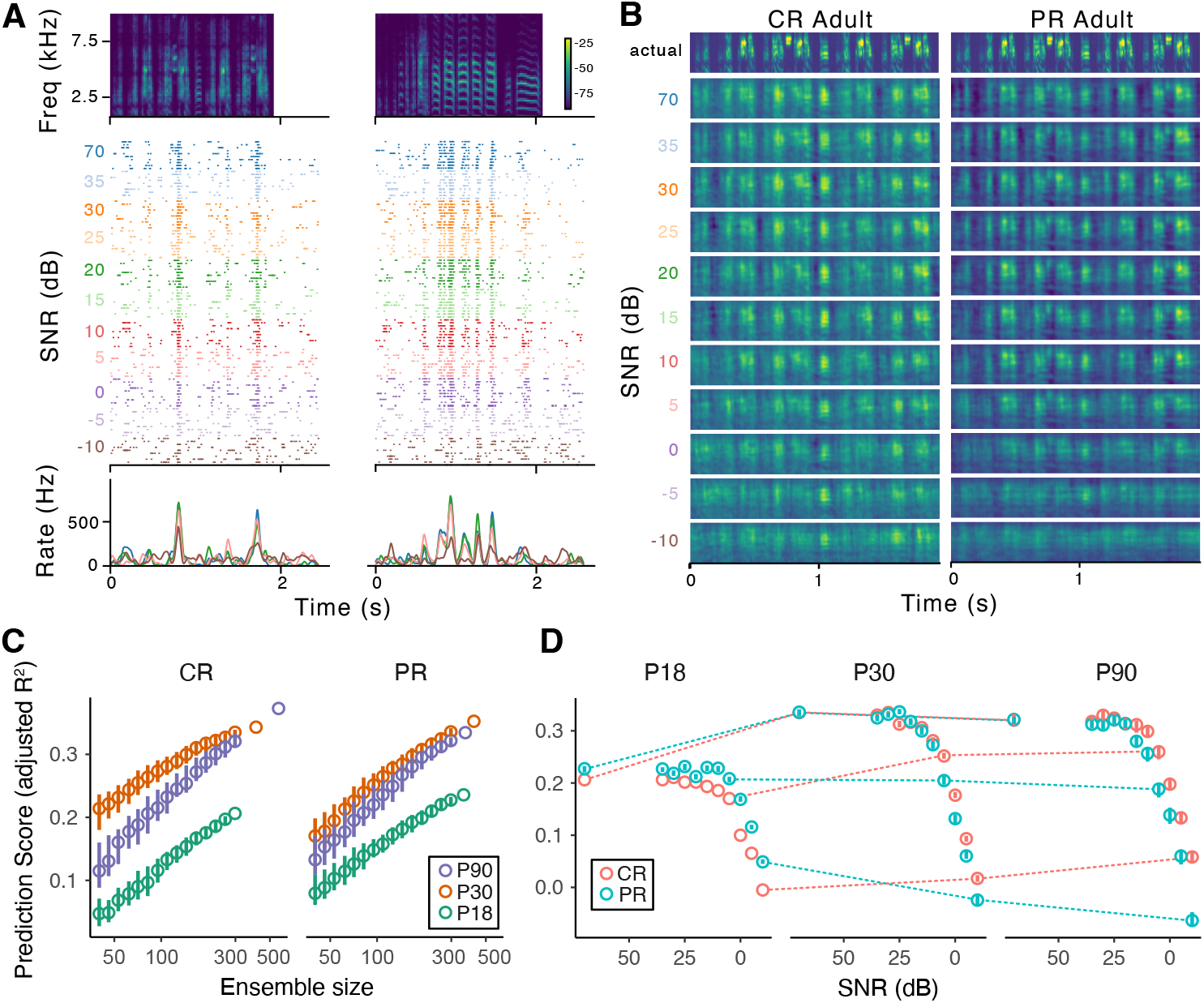
Decoding spectrographic features of song from population responses to song embedded in colony noise. (**A**) Raster plots (middle) of an exemplar neuron in response to two of the ten songs (top) at different SNR levels. Bottom plot shows smoothed firing rate histograms color coded by SNR for selected SNR levels. Spectrogram color scale is dB FS. (**B**) Example decoded test stimuli using a linear decoder trained on 9/10 songs using ridge regression (see Methods) and then tested on the held-out song. A gammatone filter bank was used for the spectrographic transform, resulting in a spectrum with log-spaced frequency bands that emphasizes lower frequencies. Top spectrogram shows the actual foreground stimulus; lower spectrograms show stimulus decoded from the response to this stimulus embedded in noise at different SNR levels. Color scale is not shown because the gammatone amplitude values are not physically meaningful. Left panels are decoded from responses of all 568 CR adult neurons and right panels from all 369 PR adult neurons. (**C**) Decoder performance on test stimuli at 70 dB SNR as a function of ensemble size. The decoder was fit using subsets of the PR or CR units at each age, sampled without replacement (*n* = 100 replicates per population size). Circles indicate median prediction score, whiskers the range between the 25% and 75% quantiles. Whiskers are not shown when the ensemble size is the same as the total number of units in that condition because there was only one replicate when this was the case. (**D**) Decoder performance as a function of SNR for randomly selected groups of 298 units from each age and rearing condition (298 was the smallest number of units in a condition). Dashed lines connect selected SNR levels across age for each rearing condition.

To control for the effect of differences in the number of units sampled from each condition, we tested decoder performance at 70 dB SNR on randomly selected subsets ranging in size on a logarithmic scale from 40 to 298 units, with 100 replicates per size (Fig. 6C). As we previously reported, decoder performance increased with the number of units included the analysis, but there were clear differences between conditions. Performance was lowest at P18 in both CR and PR birds. In the CR birds, overall performance improved between P18 and P30, and then there was an increase in the slope of the relationship between ensemble size and performance between P30 and P90. In the PR birds, the slope and overall performance increased together at P30.

Consistent with the examples (Fig. 6B), the decoded stimuli became less like the original foreground as SNR decreased (Fig. 6D). Surprisingly, at P18 the PR units had higher performance across most SNR conditions compared to units from CR birds at the same age. For the CR birds, decoder performance improved with age at every SNR. For the PR birds, performance only showed improvement at SNR levels above 5 dB and became worse at the highest levels of noise (lowest SNRs).

Overall decoder performance in adult birds was lower than what we reported previously (*R*^2^ ≈ 0.4 for comparable population sizes and stimuli), and we failed to replicate the difference between CR and PR birds at high SNRs (Moseley and Meliza, 2025). This discrepancy may reflect this study’s focus on L3 and NCM, which are more nonlinear than the other subdivisions of the auditory pallium (Sen et al., 2001; Meliza and Margoliash, 2012). To address this limitation of the linear decoder, we conducted an alternative analysis of noise invariance at the single-unit level using spike train similarity, which does not rely on identifying a linear mapping between stimulus and response. The stimuli comprised 10 songs arranged into 10 different foreground sequences, which were embedded in a fixed background of synthetic colony noise at varying SNRs (Fig. 7A). Figure 7B shows the response of a representative neuron to these auditory scenes. When responses are sorted by foreground sequence, the firing patterns remain consistent despite increasing background noise (Fig. 7B, top). Conversely, when the responses are sorted by background level (SNR), it becomes apparent that the background is driving the response regardless of the foreground once the SNR is at −5 dB or lower (Fig. 7B, bottom). For each auditory scene, we computed the similarity between the response to the scene (between 35 and −10 dB SNR) and the response to the foreground alone (70 dB SNR; Fig. 7C, top) to obtain a measure of how much the response was driven by the foreground. For each SNR level, we computed the similarity between responses to each of the foreground scenes (Fig. 7C, bottom) to obtain a measure of how much the response was driven by the background. The difference between these two averaged similarities was taken as a measure of invariance (Fig. 7D), with zero indicating that the response was driven equally by the foreground and background. s

**Figure 7.**
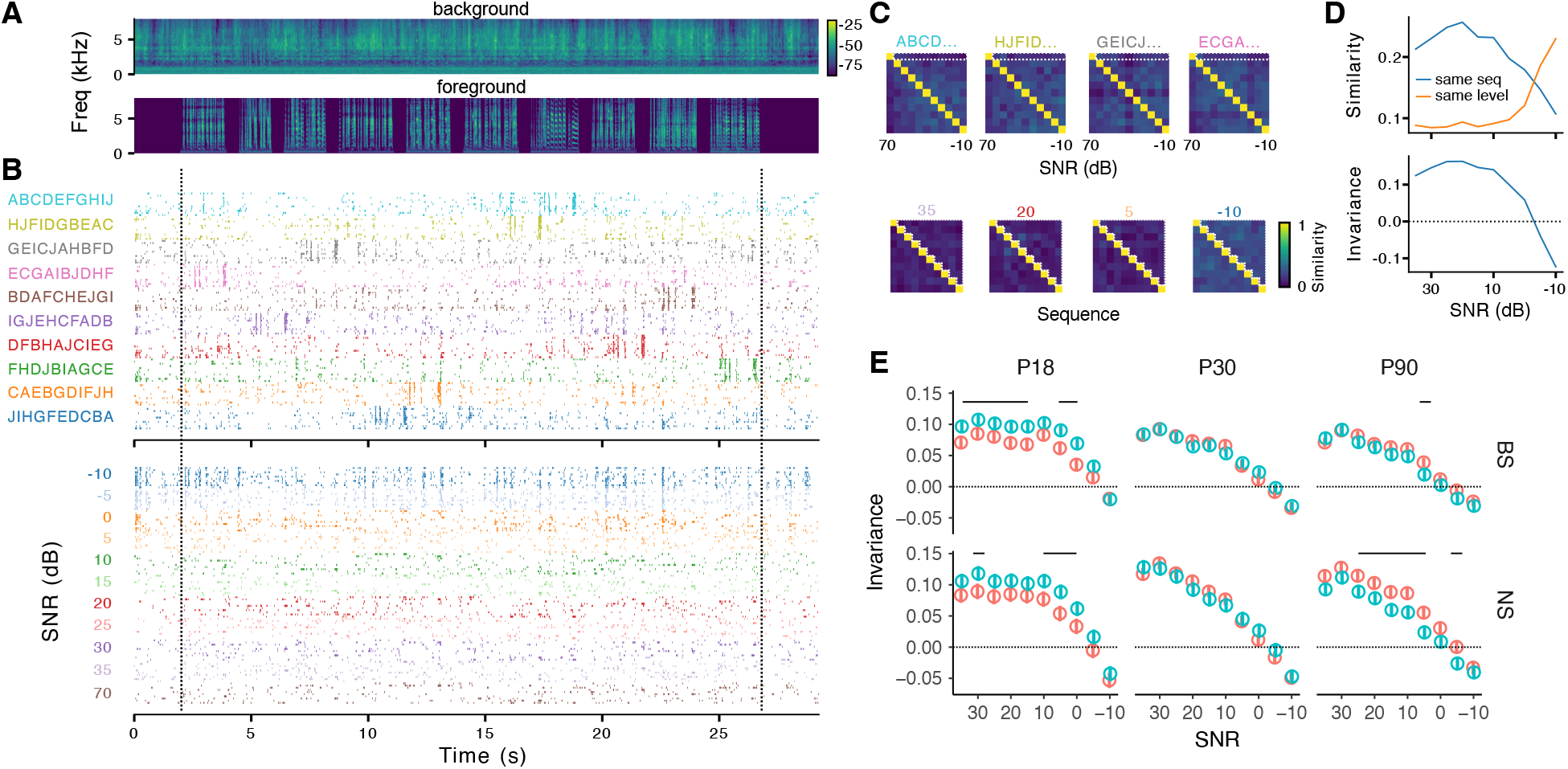
Noise invariance depends on age and experience. (**A**) Spectrograms of synthetic colony noise used as background (top) and sequence of 10 songs used as foreground (bottom). Spectrograms have the same color scale (dB FS). (**B**) Top: raster plots of an example single unit’s response to song sequences embedded in colony noise. Rasters are grouped by sequence and then by SNR (highest to lowest, going from top to bottom). Letters on the left indicate order of songs in each sequence. Bottom: raster plots of the same data, but grouped first by SNR level and then by sequence. Dashed vertical lines show beginning and end of the foreground sequences. (**C**) Quantification of invariance using spike train similarity for the example unit in *A*. Top: each trial of a given sequence was compared to the 70 dB SNR trial for that sequence (dashed white lines) to measure similarity to the foreground response. Bottom: each trial at a given level was compared to trials for all the sequences at the same level to measure similarity to the background (all sequences had the same background). (**D**) Top: for the example unit, average similarity to the corresponding 70 dB SNR sequence (blue line) decreased as the SNR decreased while average similarity to different sequences at the same level (orange line) increased. Bottom: invariance, defined as the difference between the similarities to the foreground and to the background. (**D**) Noise invariance across spike type, rearing condition, and age group. Circles represent marginal means for colony-reared (salmon) and pair-reared (cyan) conditions with 90% credible intervals for the whiskers (GLMM; see Materials and Methods). Solid horizontal bars indicate statistically significant (*p <* 0.01) differences between CR and PR at given SNR. Averaging across SNR, invariance at P18 was greater in PR birds compared to CR birds for both cell types (BS: *β* = 0.022 *±* 0.007; *z* = −3.3; *p* = 0.001; NS: *β* = 0.024 *±* 0.009; *z* = 2.68; *p* = 0.008). At P90, invariance was greater in CR birds compared to PR birds in the NS units only (*β* = 0.023 *±* 0.008; *z* = 2.88; *p* = 0.004).

Consistent with the results from the linear decoder, units from PR birds were more invariant on average than units from CR birds at P18, but this difference reversed by P90 (Fig. 7E). This reversal was due to a decrease in invariance for the PR units from P18 to P30 (BS: *β* = −0.030 *±* 0.006; *z* = −5.4; *p <* 0.001) and to P90 (BS: *β* = −0.040 *±* 0.007; *z* = −6.0; *p <* 0.001; NS: *β* = −0.031 *±* 0.009; *z* = −3.7; *p <* 0.001), whereas invariance remained stable for the CR units.

## Discussion

Vertebrate auditory systems require experience for normal maturation (Sanes and Woolley, 2011). In the zebra finch, normal experience during postnatal development comprises a complex acoustical landscape of songs and calls from many individuals (Elie and Theunissen, 2016; Moseley and Meliza, 2025). The present study demonstrates how depriving birds of exposure to this rich but noisy environment leads to deficits in secondary cortical areas that begin to manifest as early as 18 days of age and worsen into adulthood.

The response properties of single units in L3 and NCM remain stable in CR birds from immediately after fledging to adulthood, confirming previous results (Schroeder and Remage-Healey, 2021) and allaying concerns that the unavoidable use of different extracellular electrodes in adults and juveniles might have introduced a confounding variable. Spike waveforms (Fig. 1C), firing rates (Fig. 3A,B), the sampled proportions of cell types (Fig. 3C), and selectivity (Fig. 5B) all show no statistically significant differences among any of the age groups.

At the same time, CR birds show improvements in how auditory information is encoded. Discriminability progressively increases between P18 and P90 (Fig. 5A). Signal correlations, indicative of redundant coding, drop dramatically between P18 and P30 and remain low through P90 (Fig. 5C), and there is a concomitant increase in the information carried per unit about the spectrotemporal structure of song (Fig. 6C,D). These changes are consistent with a refinement of network connectivity while a homeostatic balance between excitation and inhibition is maintained.

In contrast, single-unit response properties in PR birds are unstable. We observed shifts in spike waveforms (Fig. 1C), spontaneous and evoked firing rates (Fig. 3A,B), the sampled proportions of broad-spiking and narrow-spiking units (Fig. 3C), discriminability (Fig. 5A), and selectivity (Fig. 5B). Most of the differences between age-matched PR and CR birds emerge at P30 and P90, but there are effects of rearing condition on population-level properties as early as P18 (Fig. 5C, 6C, 7E). We consider the most notable differences and our interpretations in turn.

Broad-spiking and narrow-spiking neurons are present from fledging to sexual maturity, but the BS neurons in PR birds have shallower rebounds at P30 and narrower widths at P90 (Fig. 1C). Extracellular spike waveforms reflect the dynamics of voltage-gated currents (Gold et al., 2007), so these differences may be indicative of experience-dependent intrinsic plasticity akin to what we have reported for another avian auditory area, the caudal mesopallium (CM; Chen and Meliza, 2018, 2020). In CM, experience-dependent expression of the low-threshold potassium channel K_V_1.1 produces changes in spike waveforms, phasic spiking dynamics, and a reduction in excitability. In NCM, experience-dependent effects have been seen as early as 20 days post hatch on extracellular spike waveforms (Schroeder and Remage-Healey, 2024) and on intracellular spontaneous rates (Kudo et al., 2020). The ionic currents responsible for these effects are not known, but a potassium leak current responsible for phasic spiking in adult NCM is a plausible candidate (Dagostin et al., 2015).

Our previous study in adults found much lower spontaneous and evoked firing rates in PR birds compared to CR birds, as well as a much lower probability of recording from BS neurons (Moseley and Meliza, 2025). Given the extracellular sampling bias for neurons that are active enough to be isolated during spike sorting, this difference is likely to reflect a decrease in the underlying distribution of firing rates. We see the same differences in this study after including two additional birds in the adult group. Surprisingly, at P30 it is the PR birds that exhibit higher evoked firing rates, around twofold higher compared to CR birds (Fig. 3B). This difference could reflect experience-dependent effects on intrinsic excitability (Chen and Meliza, 2020; Kudo et al., 2020) or synaptic connectivity within L3/NCM or in upstream areas such as CM. NS cells in NCM are predominantly GABAergic (Spool et al., 2021), though it is not known if GABA receptor currents are hyperpolarizing in the pallium at this age (Luo and Perkel, 2002). Inhibitory circuits and developmental shifts in chloride reversal potentials are intimately tied to experience-dependent plasticity and the closure of sensitive periods in the visual and auditory cortex of mammals (Hensch et al., 1998; Froemke et al., 2007; Yazaki-Sugiyama et al., 2009; Dorrn et al., 2010; Takesian et al., 2018). If elevated BS and NS firing rates at P30 in the PR birds reflects dysregulation of excitatory-inhibitory balance or its maturation, this could be one of the primary drivers of the abnormal auditory perception we observe in PR adults (Moseley and Meliza, 2025).

Indeed, the overall trend for PR birds is a gradual deterioration in functional response properties relative to age-matched CR birds, including lower discriminability (by P90), lower selectivity (by P30), higher population redundancy (by P18), and lower invariance (by P90). Surprisingly, on coding efficiency (Fig. 6) and invariance (Fig. 7), PR birds are slightly better than CR birds at P18 but worse at P90. This is not because the CR birds improve but because the PR birds worsen. We do not see any obvious explanation for why an impoverished acoustical environment would benefit coding efficiency and invariance immediately after fledging, but the idea that experience is necessary for maintenance has some precedent. Afferent activity is necessary to maintain established binocular segregation in the ferret LGN (Chapman, 2000), and dark-reared hamsters exhibit an initial phase of normal receptive field refinement in the superior colliculus that is subsequently lost due to dysregulation of inhibitory development (Carrasco et al., 2011). It is possible that experience-independent mechanisms also provide an initial organization for the avian auditory pallium that requires experience to consolidate and maintain.

The results presented here support our overall hypothesis that experience during the period when male birds are memorizing song has immediate effects on auditory processing that are broader than the formation of a single memory. These effects are separable from tutor song memorization, because PR and CR males both receive normal exposure to tutor song from their fathers. Although we have not verified this for our animals, acoustically isolating juveniles from all but their immediate family is common practice in studies of song learning, and male birds raised in this condition produce good imitations of their tutors (Tchernichovski and Nottebohm, 1998). The tremendous selective pressure to produce an attractive song probably provides birds with mechanisms to memorize and copy song that are robust enough to compensate for substantial deficits in central auditory processing, in which case song learning may not be a good indicator of auditory perception.

There are many differences between the social-acoustical environments of PR and CR birds, and it still remains to be seen which neural effects depend on the number and diversity of songs birds hear and which depend on interference and masking from the cocktail-party-like background. NCM remains plastic in adults (Chew et al., 1995; Thompson and Gentner, 2010; Macedo-Lima et al., 2021), and it will also be important to determine if the developmental window around P30 constitutes a critical or sensitive period, or if the effects of an impoverished acoustical environment can be rescued by introducing birds to a more complex environment (Yang and Vicario, 2015) or through behavioral training (de Villers-Sidani et al., 2010; Thomas et al., 2020) later in life. Understanding which environmental factors at what time points are necessary for the development of robust auditory perception for complex acoustic signals like song and human speech, along with the underlying mechanisms of neural plasticity, may have significance for addressing developmental language disorders that result from impairments of auditory processing.

## Author Contributions

SMM and CDM conceived and planned the experiments. SMM carried out the experiments. SMM and CDM analyzed the data, interpreted results of experiments, and prepared the figures. SMM drafted the manuscript. SMM and CDM edited and revised the manuscript and approved the final version.

